# On the relationship between afterimages and a simple sensorimotor reaction

**DOI:** 10.1101/2024.01.23.576541

**Authors:** A.A. Kulakov

## Abstract

Afterimage phenomena (afterimages) are currently widely used in a number of human activities. In this paper, the dynamic properties of afterimages are used and the comparison and relationship with the reaction to light stimulation is investigated. . It is found that the attenuation constant of afterimages and the decay constant of the variable part of the latent time of a simple sensorimotor reaction are in direct linear dependence. In addition, variable light stimulation, which contributes to the rapid attenuation of the afterimage, also accelerates the decline of the variable part of the latent time of a simple sensorimotor reaction, while the constant part increases. Based on these data, we conclude that these two phenomena, the afterimage and a simple sensorimotor reaction are interrelated in the functioning of the reflex chain of excitation transmission from sensors to the motor cortex.

## Introduction

The phenomenon of an afterimage appearing on a homogeneous background after removing an object or image from the field of view, called afterimage, is widely used in cinematography, television, and situation control systems. It has been known at least since the time of Newton and Helmholtz, but is still the object of research [1,2]. This is probably due to the complexity of the processes and structures of the brain involved in the manifestation of afterimages. There are also sound [3] and olfactory [4] afterimages.

The study of afterimages was conducted from different directions. The results showed that the brightness of the object, exposure time and background brightness were the main factors affecting the duration of the afterimage, the color difference between the stimulus and the background, and visibility [5]. At the same time, for black-and–white images, if the object was brighter than the background, then the afterimage was positive, if the background was lighter, then the afterimage was negative [5,6]. At the same time, it was found that bright afterimages caused by a decrease in brightness were stronger and lasted longer than dark afterimages created by brightness increments of equal sizes [7].

When considering chromatic afterimages, then, depending on the intensity of the background, the afterimages had a direct color, or a color of an additional color, as different as possible from the additional color of the object [8-12]. At the same time, the authors [12] did not notice the influence of age on the effect.

If we talk about the rate of disappearance of the afterimage, then different values are known here: from hundreds of milliseconds [7,13 – 15] to tens of seconds or more [5,16, 17]. The relaxation time of the afterimage is known to be 4-8 seconds [17,18]. The authors [12] did not notice the influence of age on the rate of disappearance of the afterimage. At the same time, the destruction began with the disappearance of the contour and further blurring of the afterimage [19].

The cause of the afterimages is unclear. The review [2] discusses several causes of afterimages. 1. Pigment discoloration 2. Light emission by living cells 3.Discrete dark noise 4.Biophoton radiation 5.Phosphenes 6.Delayed luminescence. Based on a critical analysis, the authors conclude that the source of afterimages, along with the discoloration of rhodopsin, may be biophoton radiation induced, for example, by glutamate. The authors [16] examined the contribution of various forms of rhodopsin and their transformations to the appearance of afterimages. The authors [20] associate afterimages and their changes with adaptation to light and changes in the content of intracellular Ca2+ concentration, in turn, regulated by the quenching of light-activated rhodopsin, acceleration of cGMP synthesis by guanylate cyclase and increased affinity of cGMP-controlled ion channels to the nucleotide, as well as the lifetime of phosphodiesterase, i.e. exclusively with events inside photoreceptor cells retinas.

The peculiarity of afterimages is their dependence not only on the brightness of the object of excitation and background, but also on other factors. Thus, the transfer of attention to another object led to the disappearance of the afterimage [1]. Also, when one eye was exposed to pulsed irritation (7-10 hz), it led to the rapid disappearance of the afterimage on the other eye [21,22].

Considerable attention was drawn by researchers to the phenomena of afterimages in connection with orientation in space. Thus, the authors [23] showed that retinotypic memory (i.e., eye-oriented memory) is less accurate than spatial memory associated with the presence of surrounding scenes. The authors [24] found that the afterimage of the hand on the table formed after a bright flash in the dark disappeared if the hand moved further in the dark. The authors [25, 26] showed that Emmert’s law, which demonstrates that the viewing distance determines the perceived size of afterimages in accordance with the number of available depth signals, could be distorted if these depth signals were absent. Zenkin spoke more definitely about this [27,28]. He believed that a structurally rich visual scene allowed the visual system to form a stable field of view and calculate the direction of the observer’s gaze only on the basis of signals in the visual stream. The hypothesis allows a coherent interpretation of the main phenomena observed in the experiments. And the observation of an oblique 3D image with the head tilted to the shoulder clearly indicated that the object of vision was not the images on the retina, but a model of the internal visual space (IVS).

These observations allowed the authors [29] to make the assumption that the Riemannian visual–somatosensory–hippocampal associative memory network can explain the perception that occurs when viewing afterimages in combination with body movement. In other words, afterimages are somehow included in the creation of an image of the surrounding space.

Currently, the location of the neurons responsible for the appearance of the afterimage is identified at various loci. For example, the authors [30] believed that afterimages can be canceled by adding real images, assuming that afterimages and real images are processed by similar mechanisms.. They came to the conclusion that the afterimages are presented similarly to their real counterparts at the level of the cerebral cortex. On the other hand, the studies of the authors [31] allowed us to say that negative afterimages cause activity in the V8 region, which is outside the area corresponding to V4 in macaques.

Thus, the authors [32], using negative color afterimages in a functional MRI study, investigated the brain mechanisms underlying changes in visual perception as a function of improving attention, which was achieved through prolonged practice of concentrated meditation. They “found increased activity in right lateralized inferior occipital and inferior frontal cortex, which suggests the importance of attentional control in modulating visual awareness.” The authors [25] suggested that the neural mechanism underlying both the tilt illusion and the tilt aftereffect includes inhibition tuned to orientation in V1. The authors [7] associated afterimages with the dorsal lateral geniculate nucleus (dLGN) and the primary visual cortex (V1) of cats and monkeys. The authors [33] associated the appearance of phosphenes with the V1 region. On the other hand, the authors’ studies (31) allowed them to say that negative afterimages cause activity in the V8 region, which is outside the area corresponding to V4 in macaques.

Thus, it seems that the afterimages are associated with a temporary memory that allows you to compare changes coming from visual sensors. This creates significant memory savings in the central nervous system.

We were interested in another feature of afterimages, namely the gradual fading in time. This is due to the fact that we have shown that such a well-known phenomenon as a simple sensorimotor reaction, in particular, the latent time of the visual-motor reaction is a variable and, being significant immediately after responding to irritation, then gradually decreases after some time to a constant value [34,35]. Therefore, we had the idea of comparing the destruction curve of the afterimage and the change in latent time after the reaction. This comparison is the subject of this publication. In our opinion, this is the first attempt to compare together the phenomena of afterimage and sensorimotor reaction.

## Methods

The research involved 22 people of different ages and genders. All the subjects are volunteers, colleagues of the author, including himself. All studies were conducted in the spring of 2019, before the pandemic. In addition, in 2022-2023, volunteer students participated in part of the research. Since we have shown that for even a few minutes the parameters of a simple sensorimotor reaction are subject to fluctuations [35], we tried to carry out measurements as quickly as possible (1 series of measurements at a time) or repeated 1-2 series of measurements for several days.

### Determination of the parameters of a simple sensorimotor reaction

The parameters of a simple sensorimotor reaction were determined in the same way as in [34]. In short, it consisted of the following. A red square spot of 5 mm in size appeared on the dark computer screen. The subject, in response to this, pressed the mouse button and thus extinguished the spot. In response to this, after waiting, a new spot appeared, and a new button press followed. In total, there were 50 spot presentations in the series. There were no breaks in the series, so the waiting time for the n-spot was calculated from the moment of reaction to the n-1 spot of the series until the appearance of the n-spot, and the latent time – from the moment of the appearance of the spot to pressing the button in response to this appearance. The waiting time was presented in accordance with a random counter of the BCB++ 6 program, which controlled the measurement process. All results were recorded and then calculations were performed. At the first stage, the waiting times were ranked in ascending order and the latent times were placed in a row corresponding to them. Then the values of latent times that were too much different in the direction of increasing for large waiting values were either thrown out or replaced with averages compared to neighboring points. Similarly, with small waiting times, too short latency times were either thrown out or replaced with average ones compared to neighboring points. Usually the number of such points was no more than 5-10%. Then the latency times were ranked from large to smaller. As shown by Kukinov [36], this approach is mathematically rigorous and reduces the average error. This decrease in values is explained by the fact that a variation series of the error distribution is superimposed on the monotonic decline in latent time, which also decreases monotonically when we rank our data. At the next stage, at the end of long waiting times, the inflection value was selected, from which a sharp decline in values began. Usually this point was about 15-20% from the end. From this point, the subsequent ones were fixed at the same level. This is quite simply explained by the fact that we are dealing with a monotonically decreasing curve, which from about two-thirds of the dependence goes to the stationary. We have previously verified this by summing up several curves obtained without processing. The resulting difference between the ranked values and the fixed value was symmetrically subtracted from their first 15-20% points from the beginning of the dependence. In some cases, repeated ranking was done in descending order because, due to the relatively small number of points, dips could appear with symmetrical subtraction. It was as if we rotated the ranked dependence counterclockwise, taking the midpoint as the center, while most of the error variation series was subtracted. This is illustrated in Figure 1.

**Fig. 1.**
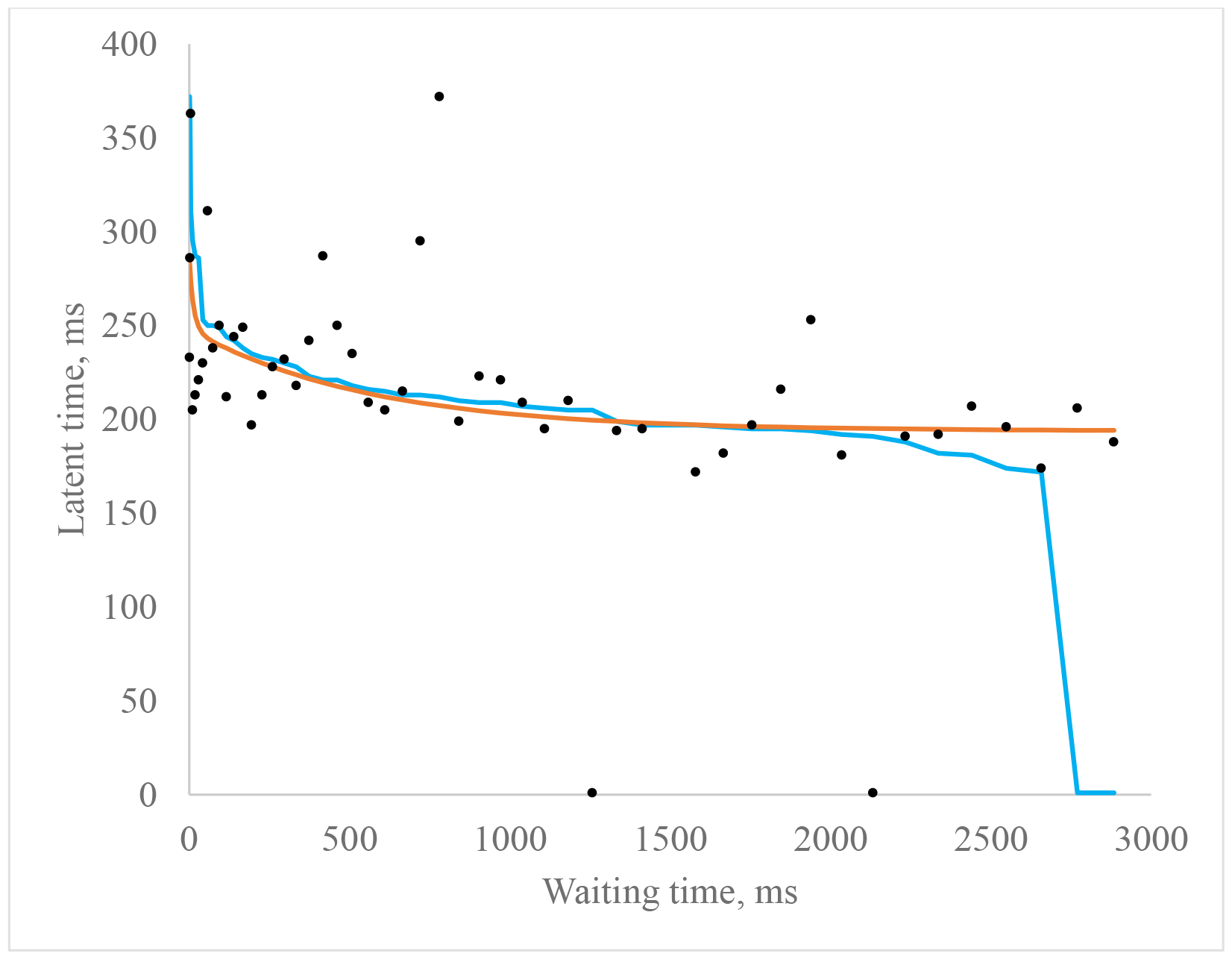
The course of the “reconstruction” of the effect of the dependence of the latent time of SSR on the waiting time. Black circles are the experimental data. Blue line is the experimental data ranked by reduction. Red line is the ranked data minus the error variation series.

After that, using the “solution search” program in the ExCel 2010 package, multi-exponential regression was found using the least squares method, based on the exponential decline model according to the formula

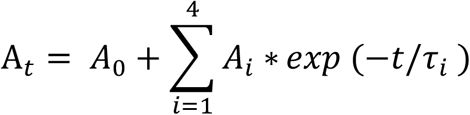

where A_t_ is the theoretical value of the latent time during t, A_t_ is the theoretical value of the latent time at the hospital, i is the number of components, A_i_ is the i component at t=0, τ_i is the relaxation time for component i. A given amount of i=4 was given in excess, usually 2-3 components were obtained, the rest – with zero amplitudes.

### Decay of the afterimage

Measurements of attenuation after phases were carried out on the installation using an Arduino Uno microcontroller. The light stimulus was white LEDs placed in cubic illuminators with an output square window of 18 mm in size . A system of reflectors reflected light from an LED onto a window covered with a matte plastic film and translucent paper, so that the entire square window was illuminated evenly.

Both illuminators were located side by side. They were placed in a darkened box with a cutout for the eyes opposite, at a distance of 20 cm from the cutout. In one window the brightness was 1.52 cd, in the other it varied from 0.067 cd to 0 cd. Brightness changes were carried out using a luxmeter-yarkometer “TKA-PKM” (NTP “TKA” SPb). The illuminators were supplied with voltage from the Arduino PWM outputs. The frequency of voltage supply to the PWM outputs is 976 Hz. When changing the signal range set by Arduino, the average brightness can be changed within wide limits. After a 5-minute adjustment to the darkness, the LED in the right illuminator turned on for 2 seconds, and then turned off. The LED turned on immediately on the left illuminator, so that the left window shone with different average brightness. The subject compared the illuminated left window with the apparent brightness on the right window, and at the moment of apparent brightness equality pressed the button. The number of experiments in the series was 40, with 4 representations of each brightness, including darkness.. There were several such series, one or two per day, and all the data were averaged. Thus, the time dependence of the apparent brightness of the afterimage on the time after switching off the exciting light was obtained. It should be noted that some of the subjects could not notice the afterimages at all, for some they had to make a training program for mastering, this is due to the small number of subjects.

## Results

In Fig. 4 shows the dependence of the relaxation time of the slow component of the variable latent time on the relaxation time of the afterimage. The linear relationship between these two processes is quite clearly manifested: the attenuation of the afterimage and the attenuation of variable latent time. The correlation coefficient is 0.923. The Kendall coefficient is 0.71 with a significance of 0.005. Both of these values show that there is a direct linear relationship between these two changes, so we concluded that both of these processes are interrelated.

Based on this fact, it can be assumed that in the attenuation of the afterimage and the latent time of a simple sensorimotor reaction, there is a functioning of a common mechanism.

Consequently, factors affecting the manifestation of the afterimage can also affect the SSR. For example, it was shown that irritation of one eye with variable (7-10 hz) light suppressed the manifestation of an afterimage [21,22]. We used this technique. We assumed that the components of the decline of the variable part of the PSMR will either disappear completely or accelerate, which can be explained as the disappearance or decrease of the variable part of the SSR.

It should be noted that this dependence does not always manifest itself. When measured with one eye (approximately 30-40% of the subjects), there was practically no decline, or there was, but at first an increase in latency developed, and then a decrease. In addition, the variability of latent time was greatly increased.

Table 1 shows the effect of alternating light (10 hz, i.e. 50 msec light, 50 msec dark) on the parameters of the SSR. As can be seen from the table, irritation of one eye with variable illumination on a dark background led to a significant decrease in the relaxation constants of the variable part of the latent time, with the exception of one case when the relaxation constant of the slow component, on the contrary, slowed down.

**Table 1.** The effect of irritation of the left eye with variable light on the parameters of the SSR of the right eye on a dark background.

| Subject<br>(gender) (age) | Impact | A<sub>1</sub>, ms | T<sub>1</sub>, ms | A<sub>2</sub>, ms | T<sub>2</sub>, ms | A<sub>0</sub>, ms | Average<br>error |
| --- | --- | --- | --- | --- | --- | --- | --- |
| M. (w) (22) | Control | 159,5 | 198,4 | 39,3 | 1754,6 | 254,0 | 3 |
|  | Irritation of eye | 134,3 | 27,7 | 107,3 | 862,3 | 281,6 | 3,9 |
| K. (w) (23) | Control | 218,1 | 84,0 | 55,0 | 846,4 | 315,6 | 4,3 |
|  | Irritation of eye | 2803,5 | 4,6 | 180,9 | 292,1 | 332,6 | 8,8 |
| B. (m) (23_) | Control | 138,4 | 68,9 | 38,3 | 723,1 | 308,2 | 2,0 |
|  | Irritation of eye | 68,6 | 34,8 | 98,4 | 687,7 | 307,7 | 5,1 |
| N. (m) (23) | Control | 133,3 | 106,2 | 52,6 | 1262,4 | 330,7 | 5,4 |
|  | Irritation of eye | 213,7 | 37,4 | 185,4 | 702,5 | 365,8 | 9,5 |
| C. (m) (35) | Control | 232,0 | 96,7 | 29,7 | 2753,4 | 238,9 | 6,0 |
|  | Irritation of eye | 54,7 | 28,1 | 85,2 | 427,6 | 280,6 | 3,6 |
| P. (m) (25) | Control | 128,8 | 121,1 | 33,4 | 1809,2 | 238,4 | 4,2 |
|  | Irritation of eye | 155,3 | 13,1 | 149,8 | 1443,0 | 333,6 | 5,3 |
| G. (w) (35) | Control | 87,8 | 27,1 | 47,5 | 771,9 | 279,8 | 1,3 |
|  | Irritation of eye | 44,9 | 17,4 | 88,9 | 968,8 | 331,4 | 2,4 |
| D. (ж) (11) | Control | 54,3 | 81,9 | 49,2 | 2598,9 | 338,7 | 2,8 |
|  | Irritation of eye | 88,6 | 14,6 | 121,4 | 626,2 | 444,3 | 4,6 |
| B. (w) (58) | Control | 111,5 | 172,3 | 61,4 | 1606,3 | 264,0 | 6,3 |
|  | Irritation of eye | 141,22 | 112,2 | 66,7 | 948,9 | 288,8 | 4,1 |
| B. (m) (70) | Control | 25,6 | 164,6 | 74,2 | 1561,0 | 240,6 | 2,5 |
|  | Irritation of eye | 312,3 | 0,99 | 69,9 | 795,4 | 281,9 | 2,1 |

However, in all cases, the value of the constant component of the latent time increased significantly. Thus, from the totality of the results, with irritation of the left eye, compared with the control without irritation, the relaxation constants of the decay of the variable part of the latent time decreased, while the constant part increased significantly.

It should be noted that we usually determined the parameters of the PSMR on a computer when there was a sufficiently illuminated background, whereas under the conditions of measuring the afterimage, and then variable irritation, it was preferable to carry out in the dark. Therefore, in the next study, we measured the SSR on a computer by placing a flashing white LED with the same flashing frequency in the field of view of the left eye.

The study data are presented in Table 2. As can be seen in the table, irritation with white pulsed light dramatically accelerated the relaxation time of the latency decline, as it was on a dark background. The action was similar to the relatively fast and slow component. At the same time, the constant part of the latent time on a dark background significantly increased, and on a light background this decrease was insignificant.

**Table 2.** The effect of intermittent light irritation of the left eye on the parameters of the right eye SSR on a light background.

| Subject<br>(gender) (age) | Impact | A<sub>1</sub>,ms | T<sub>1</sub>,ms | A<sub>2</sub>,ms | T<sub>2</sub>,ms | A<sub>0</sub>,ms | Average<br>error |
| --- | --- | --- | --- | --- | --- | --- | --- |
| A. (w) (23) | Control | 141,1 | 125,3 | 59,2 | 742,7 | 213,9 | 7,25 |
|  | Irritation of eye | 77,2 | 44,8 | 128,6 | 484,5 | 232,8 | 5,36 |
| G. (w) (22) | Control | 124,3 | 31,8 | 132,7 | 1204,1 | 214,2 | 5,11 |
|  | Irritation of eye | 74,6 | 8,0 | 133,8 | 669,5 | 235,0 | 5,52 |
| D. (w) (22) | Control | 208,6 | 18,2 | 226,3 | 1195,0 | 312,9 | 16,8 |
|  | Irritation of eye | 752,6 | 2,1 | 1082,1 | 460,5 | 498,3 | 36,34 |
| I. (m) (23) | Control | 86,4 | 15,8 | 123,7 | 479,8 | 236,6 | 5,14 |
|  | Irritation of eye | 36,5 | 0,03 | 127,9 | 273,4 | 242,7 | 6,62 |
| P. (m) (23) | Control | 124,8 | 41,6 | 78,8 | 748,9 | 228,3 | 4,96 |
|  | Irritation of eye | 489,8 | 0,5 | 108,9 | 625,5 | 217,0 | 4,64 |
| Sh. (m) (23) | Control | 67,4 | 7,7 | 159,5 | 771,5 | 246,0 | 9,21 |
|  | Irritation of eye | 72,8 | 7,53 | 46,6 | 423,2 | 254,4 | 2,61 |

Next, we changed the measurements so that the determination of the latent time of the right eye was carried out with a waiting time of 5000 ms, i.e. at the level of constant latent time. At the same time, the frequency of the light irritating the left eye was changed. We can see this in the diagram in Fig. 5. With the exception of frequencies 20 and 16.6 hz, all other latent time values significantly exceeded the latent time without illumination of the left eye at the level of p<0.05 for frequencies 25 and 12.5 hz, and p<0.01 for other frequencies. Note that with constant illumination of the left eye (N), the increase in latency compared to darkness (T) is unreliable.

Figure 5 shows data for one subject. For other subjects, the results are similar, only the frequency with the largest amount of latent time and the absolute value of the latter can change. Thus, it was shown that irritation with pulsed light of one eye has a significant effect on the measurement of the parameters of the SSR of the other eye.

## Discussion

The results presented by us suggest that the phenomenon of afterimage is somehow related to the functioning of a simple visual-motor reaction. This is indicated by a) a linear relationship between the relaxation times of the afterimage and the latent time of SSR, b) the effect of pulsed excitation of one eye on the afterimage and on the parameters of the SSR.

At the same time, statement a) reliably concerns the slow component of relaxation, since we did not capture the fast component accurately enough. However, as we showed in [34], there is a direct correlation between the fast and slow components, and we can leave this statement in force for now. The phenomenon of the difference in the behavior of slow relaxation is similar to the manifestation of an afterimage on different backgrounds: the afterimage from light irritation is seen on a dark background as light, and on a light background as dark. A similar pattern is observed when stimulated by chromatic light – in one case, the afterimage has a direct color, in the other – a color in an additional color [8-12].

Nevertheless, the ability of pulsed light stimulation to accelerate the relaxation of the kinetic parameters of the SSR is similar to the effect of this irritation on the acceleration of the decline of the afterimage [21,22]. Thus, the previously mentioned similar behaviors of afterimages and latent time allow us to talk about a common mechanism that affects both.

At the same time, in addition to the accelerating effect of impulse stimulation, there is also an increase in constant latency time. The latent time of the right eye is significantly increased, although only the left eye is irritated. At the same time, the greatest increase in latency is observed at low frequencies of irritation (0.5-2 hz). Recently, a number of articles have appeared on multimodal interaction in the stimulation of different receptors. In this case, we are dealing with identical, paired receptors. Apparently, this inhibitory effect can take place before the visual cortex.

We can draw a similar conclusion from our preliminary report, where alternating light irritation of one eye, then the other, and SSR in response were studied [38,39]. There, as we expected, the relaxation of latent time decreased, since during the reaction of one eye, the other restored its ability to react at that time. The same applied to the audio signal and the response with different fingers. However, this acceleration of relaxation was much less than could be expected, because even if the response was missed, even the shortest waiting time, the total cycle time was: throughput waiting time, latency throughput time, waiting time to respond to, i.e. the total waiting time was already on the order of 500-1000 ms instead of 1-300 ms. This is also explained by the fact that at the places where the signal from paired receptors is combined, there is a place to choose between one and another signal. In other words, a simple sensorimotor reaction actually involves choosing between two signals.

Thus, the phenomenon of afterimage is associated with the manifestation of a delay when the eye is irritated by light. Here it is necessary to turn to the research of afterimages. We said above that afterimages are part of creating an image of the surrounding space, or in general, the surrounding world, moreover, according to [29], the location of the afterimage is a distributed visual–somatosensory-hippocampal associative network, and all this is intended for the most unambiguous awareness of the subject in space. As a clarification of the working hypothesis, we propose the following. During the exposure of light to the eye, a certain fund of excited neurons is formed, which exists for some time and gradually disappears in the dark. From Fig. 2,3 we could estimate the size of this fund. The maximum value of the apparent brightness of the afterimage, about 48 conventional units, was comparable to the maximum brightness in the left window, i.e. 0.067 cd, which corresponded to about 0.044% of the brightness of the right window in the installation. Therefore, it could be assumed that the fund of excited neurons responsible for the manifestation of the afterimage was slightly less than 5% of those neurons responsible for the perception of light, causing both an afterimage and a sensorimotor reaction. For a while, it sends excitation signals further, which is perceived as an afterimage, i.e. an apparent glow. However, he simultaneously stops perceiving arousal from visual sensors, which leads to an increase in latency time until this fund decreases to a minimum threshold. Perhaps we are dealing with the circulation of excitation at one of the stages of the reflex arc, perhaps this is the activation of inhibitory neurons with subsequent dissipation of inhibition. It is interesting to consider the observations of the authors in this regard [40]. They studied the differentiated visual-motor reaction “Go/No Go” triggered by a test stimulus (TC = 0.1 ms, Blue) against the background of a masker stimulus (MS = 30 ms, Green). It was shown in the work that in the interval between the test signal and the masking signal in the range of 30-100 msec, an increase in the latent reaction time is observed with an extension of this interval. The authors interpret this inhibition within the framework of Ivanitsky’s paradigm [41] about the cycle of excitation from the primary visual cortex to the frontal lobe of the cerebral cortex. It seems to us that here we can talk about exactly the same model that we propose, despite the measurements of the differentiated visual-motor reaction in the work [39], only we have this cycle a little earlier, and assume that there is a moment of choice for a simple visual-motor reaction. The physiological meaning of such dynamic memory is to save space and time, when the basic information about the surrounding world remains unchanged and only a small variable part is considered and analyzed – in terms of coordinates or brightness.

**Fig. 2.**
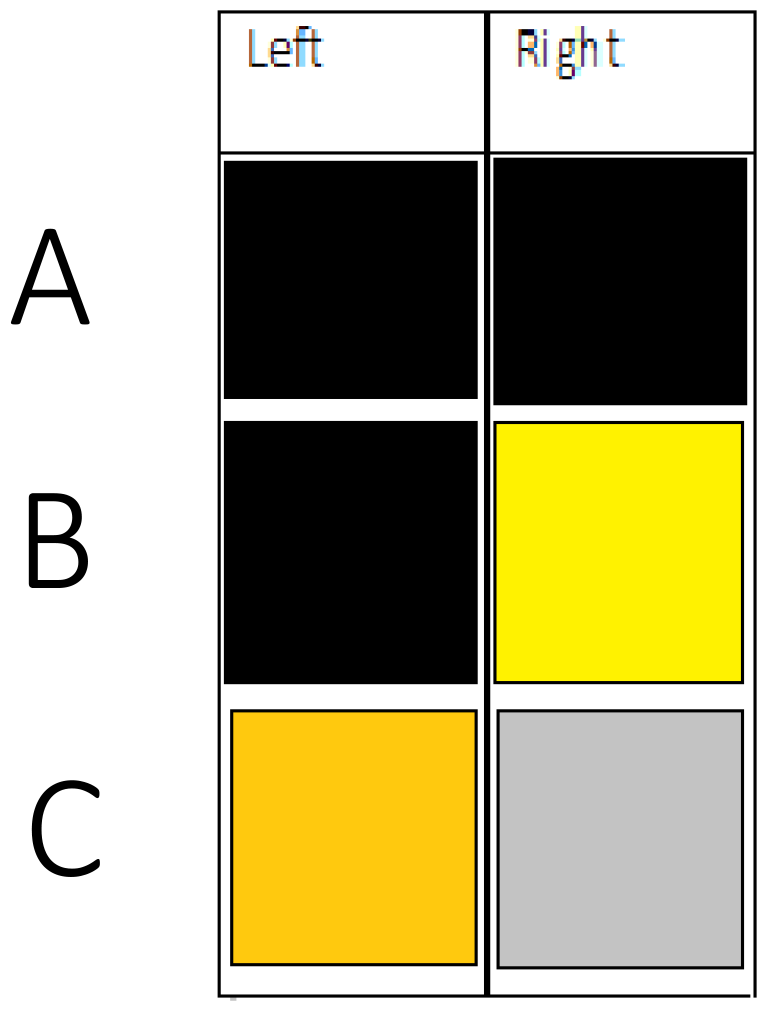
Stages of measurement after the image. A - In the Left window, and in the Right window -2 seconds of darkness, B. In the Left window –darkness, in the Right window - 2 seconds of light, C. In the Left window weak light, in the Right window – afterimage. With the apparent equality of brightness in the Left and Right windows - pressing the button and switching to the A state

**Fig. 3.**
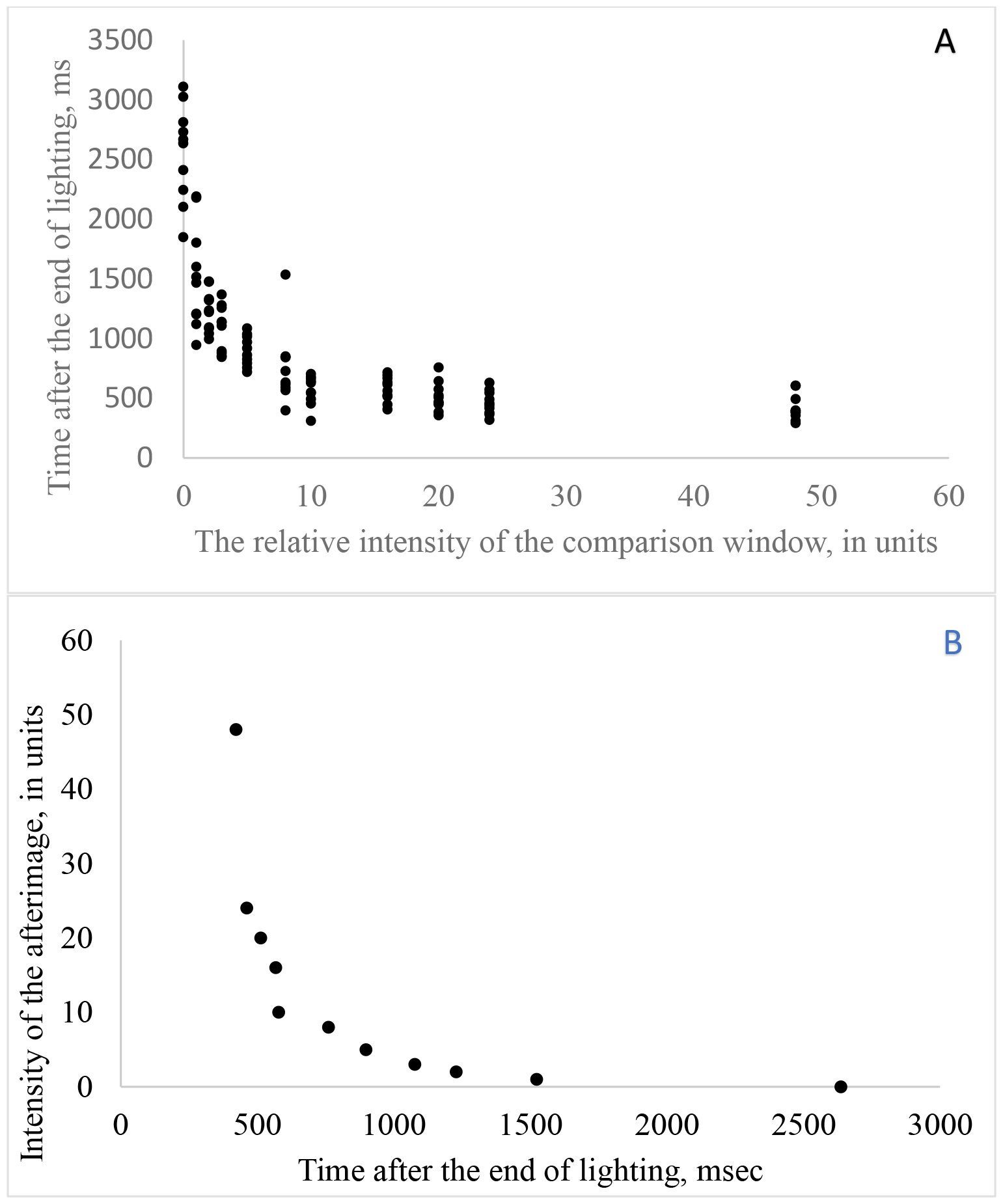
Determination of the dependence of the intensity of the afterimage caused by a 2-second pulse of light. A-experimental points of intensity of the afterimage, with equality with the intensity of the control window. B - the averaged values of the intensity of the afterimage depending on the waiting time after the light pulse.

**Fig. 4.**
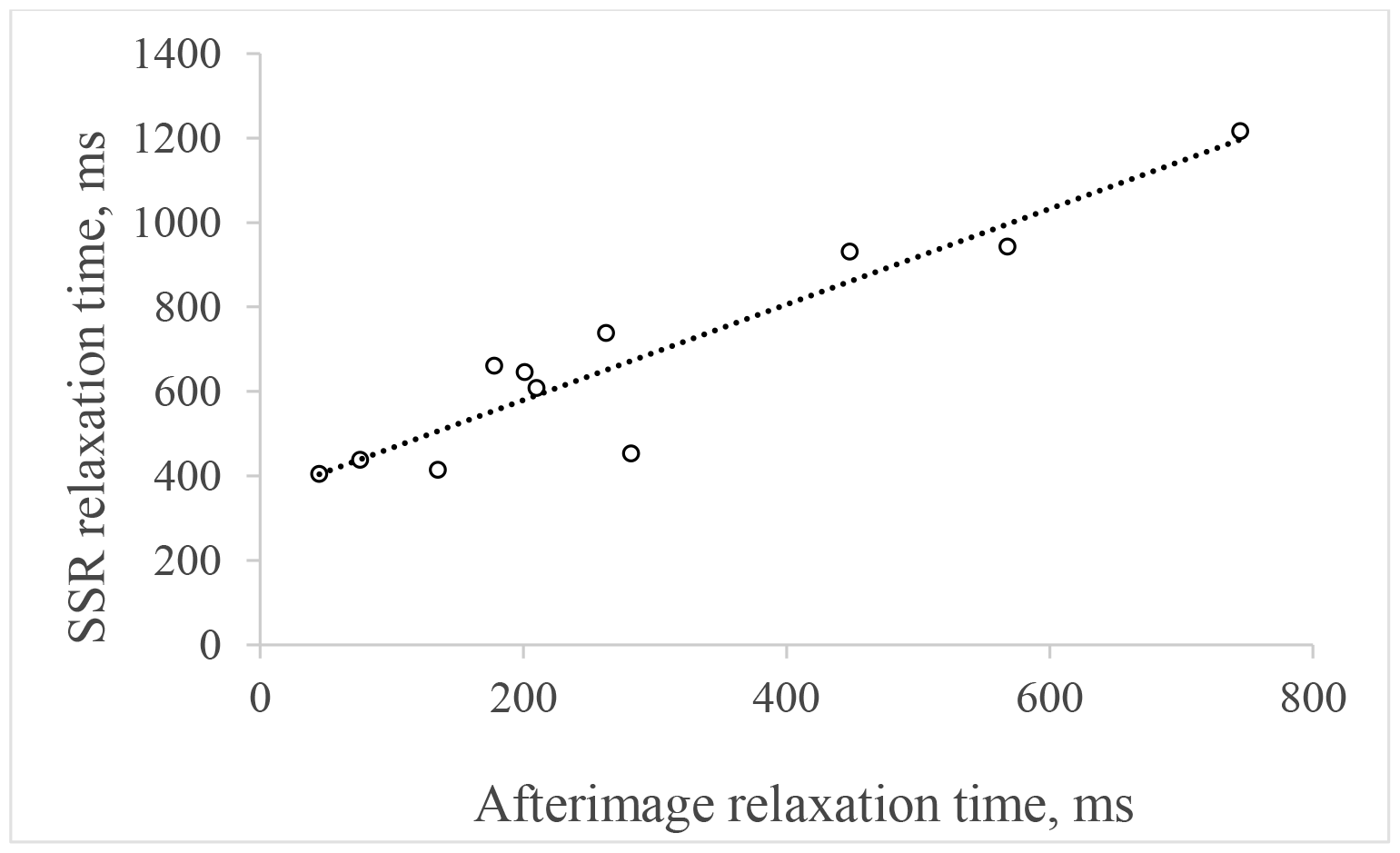
The interdependence between the relaxation time of the slow component of the latent period and the relaxation time of the attenuation of the afterimage.

**Fig. 5.**
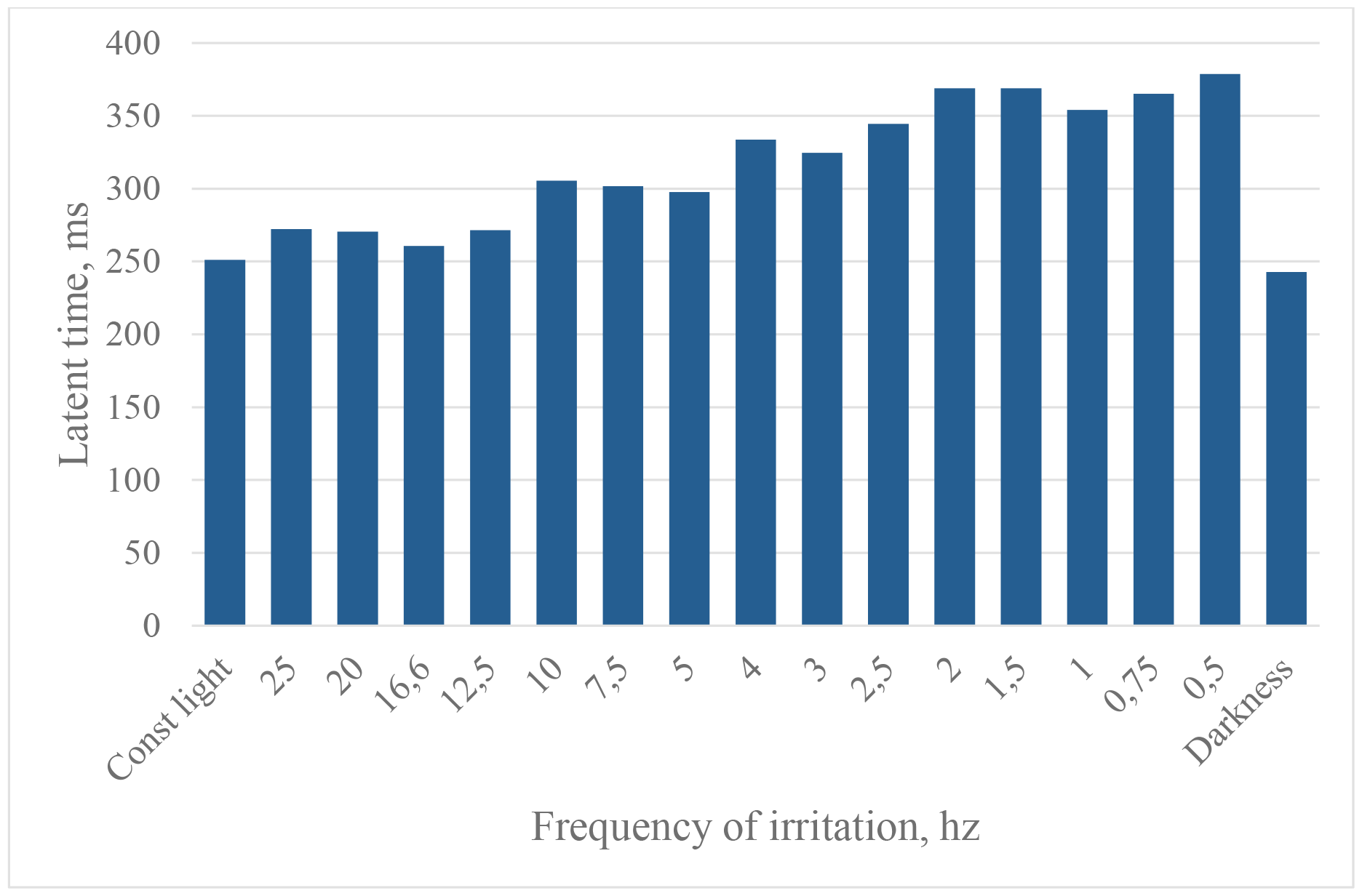
The effect of irritation of the left eye by pulsed illumination on the latent time of a simple sensorimotor reaction of the right eye. The irritation was carried out on a light background with flashes of white color (the dark interval is equal to the light one). The latency time was determined after a flash of white light with a waiting time interval of 5 seconds. On the abscissa axis is the frequency of flashes in kc, the first column is constant illumination, the last column is darkness. On the ordinate axis is the value of the latent time in msec.

Concluding our discussion, we would like to draw attention to the fact that the joint study of afterimages and a simple visual-motor reaction can significantly advance our understanding of the perception of a visual signal and the mechanism of its implementation.

Ethical standards. All studies were conducted in accordance with the principles of biomedical ethics and approved by the Local Ethics Committee of Kazan State Medical University, Kazan.

## Abbreviations

SSR: simple sensorimotor reaction

## Notes

### Competing Interest Statement

The authors have declared no competing interest.

### Summary of Updates

In comparison with the original version, the abbreviations and the name of the last figure, as well as links to it, have been corrected

